# Cyclooxygenase-dependent mechanisms mediate in part the anti-dilatory effects of perivascular adipose tissue in uterine arteries from pregnant rats

**DOI:** 10.1101/2021.01.16.426949

**Authors:** Oluwatobiloba Osikoya, Spencer C. Cushen, Styliani Goulopoulou

## Abstract

Uterine perivascular adipose tissue (PVAT) contributes to uterine blood flow regulation in pregnancy, at least in part, due to its effects on uterine artery reactivity. Here, we investigated the effects of uterine PVAT on endothelium-dependent pathways involved in relaxation of main uterine arteries. We hypothesized that uterine PVAT modulates the balance between the contribution of nitric oxide synthase (NOS)- and cyclooxygenase (COX)-dependent pathways to acetylcholine (ACh)-induced relaxation in isolated uterine arteries. Concentration-response curves to ACh (1 nM – 30 µM) were performed on main uterine arteries from pregnant and non-pregnant rats. Arteries were exposed to Krebs-Henseleit solution (control) or PVAT-conditioned media (PVATmedia) in the presence of the following inhibitors: L-NAME (100 µM), indomethacin (COX inhibitor, 10 µM), SC560 (selective COX-1 inhibitor, 1 µM), NS398 (selective COX-2 inhibitor, 1 µM), SQ 29,548 (selective thromboxane receptor (TP) inhibitor, 1 µM). Indomethacin suppressed ACh-induced relaxation in control uterine arteries from pregnant rats (p<0.0001) but not in non-pregnant rats (p>1.0). In arteries incubated with PVATmedia, the presence of indomethacin increased ACh-induced relaxation, reversing the anti-dilatory effect of PVATmedia. NOS inhibition reduced ACh-induced relaxation in uterine arteries from pregnant rats, and exposure to PVATmedia did not change this effect. Selective inhibition of COX-1 but not COX-2 suppressed relaxation responses to ACh in control arteries. The presence of PVATmedia abolished the effect COX-1 inhibition. Incubation of uterine arteries from pregnant rats with PVATmedia increased production of thromboxane B_2_ (TxB_2_, p=0.01). TP inhibition did not have any effect on the anti-dilatory properties of PVATmedia. In conclusion, uterine PVAT releases transferable factors that reduce relaxation responses to ACh via a COX-dependent mechanism in isolated uterine arteries from pregnant rats.

## Introduction

Perivascular adipose tissue (PVAT) is a metabolically active and physiologically relevant tissue that contributes to the maintenance of vascular homeostasis by modulating vascular tone (1, 2, 4, 7, 35, 41). The effects of PVAT on its adjacent vascular wall are either due to uptake and metabolism of vasoactive amines from nerves innervating blood vessels by PVAT (57) or by direct action of PVAT-derived factors (4, 18, 31) on vascular smooth muscle and endothelial cells (3, 44). The vasoactive properties of PVAT are vascular bed-specific and vary with physiological or pathological states and experimental vascular preparations (18, 31, 41, 44).

Removal of uterine PVAT from one of the two uterine arteries resulted in a 40% reduction of uterine artery blood flow (UtBF) as compared to the adjacent uterine artery with intact PVAT in anesthetized healthy pregnant rats on gestational day 16 (41). The reduction in UtBF following removal of PVAT was not observed in non-pregnant rats (41), suggesting that the interaction between PVAT and uterine artery is pregnancy-specific. In vascular preparations of isolated uterine arteries from pregnant rats, PVAT suppressed uterine artery relaxation to acetylcholine (ACh), and this anti-dilatory effect was mediated by soluble factors secreted by uterine PVAT (41). These previous *in vivo* studies demonstrated the functional role of PVAT in the uterine vasculature in rat pregnancy, while the *in vitro* studies interrogated the effects of PVAT on the uterine vascular wall in the absence of systemic and hemodynamic factors. In the present study, we sought to extend the findings from these studies investigating the mechanisms by which PVAT affects ACh-induced relaxation in isolated uterine arteries.

PVAT-derived factors modulate vascular tone by directly targeting vascular smooth muscle cells or by altering the nitric oxide synthase (NOS)/nitric oxide (NO) system in the vascular endothelium (3, 44, 49). The influences of PVAT on the NO/NOS pathway in the uterine vasculature have not been studied. Nevertheless, it is known that the endothelium is required for the anti-dilatory effects of uterine PVAT on rat uterine arteries as there were no differences in relaxation responses to sodium nitroprusside between PVAT-intact and PVAT-denuded arteries from pregnant rats (41).

In addition to NOS/NO pathway, vascular smooth muscle cell relaxation is also facilitated by endothelium-derived prostacyclin (PGI_2_) and non-NO, non-PGI_2_-related pathways, such as endothelium-derived hyperpolarization (EDH) (17, 61)(20, 29). The relative contributions of NO, PGI_2_, and EDH to ACh-induced relaxation are vascular bed specific and vary with vessel size and function (i.e., conduit vs. resistance arteries) (53, 55). During pregnancy, the contribution of EDH is predominant in radial uterine arteries, which are small resistance vessels (19), while synthesis and release of NO play a predominant role in ACh-induced relaxation in larger uterine arteries (25, 38)}. Pregnancy also upregulates vascular production of PGI_2_. For example, studies in ovine uterine artery endothelial cells showed that production of PGI_2_ increased during pregnancy as did the protein content of cyclooxygenase (COX) enzymes responsible for this production (21, 24, 32). Whether uterine PVAT mediates its effects on uterine artery dilatory responses to ACh by modulating the contribution of endothelium-dependent pathways is currently unknown.

In the present study, we investigated the effects of uterine PVAT on endothelium-dependent pathways involved in rat main uterine arteries. We hypothesized that uterine PVAT modulates the balance between the contribution of NOS- and COX-dependent pathways to ACh-induced relaxation in isolated uterine arteries from pregnant rats.

## Methods

### Chemicals and Reagents

Acetylcholine (ACh, cat#: A6625), indomethacin (cat#: I8280), N_ω_-Nitro-L-arginine methyl ester hydrochloride (L-NAME, cat#: N5751), NS398 COX-2 inhibitor (cat#: 349254), phenylephrine (PE, cat#: P6126), phenymethysulfonyl fluoride (PMSF, cat# P7626), sodium orthovanadate (Na_3_VO_4_, cat#: 450243), and SC560 COX-1 inhibitor (cat#: 565610) were obtained from Millipore-Sigma (Millipore Sigma, Saint Louis, MO, USA). SQ 29,548 (cat#: 1025) was obtained from Cayman (Cayman Chemical, Ann Arbor, MI). Stock solutions of indomethacin (10 mM), NS398 (1 mM) SC560 (1 mM), and SQ 29,548 (1 mM) were prepared in dimethyl sulfoxide (DMSO). Stock solutions of L-NAME (100 mM), PE (100 mM) and ACh (100 mM) were diluted in Milli-Q distilled water (Millipore Sigma, Burlington, MA, USA). For myograph concentration-response curves, all dilutions were prepared on the day of the experiments. T-PER protein extraction reagent (cat#: 78510), cOmplete™, mini, EDTA-free Protease Inhibitor Cocktail tablets (cat# 11836170001, Roche, Basel, Switzerland), Pierce™ bicinchoninic acid (BCA) Protein Assay reagents (Pierce™ Bovine Serum Albumin Standard, cat# 23210; Pierce™ BCA Protein Assay Reagent A, cat#: 23228 and Pierce™ BCA Protein Assay Reagent B, cat#: 23224) were obtained from Thermo Fisher Scientific (Waltham, MA, USA).

### Animals

All protocols were performed in accordance with the Guide for the Care and Use of Laboratory Animals of the National Institutes of Health and the ARRIVE guidelines (26, 27). Experimental protocols were approved by the Institutional Animal Care and Use Committee of the University of North Texas Health Science Center (IACUC#: 2017-0042).

Male (body weight and age at arrival in animal facilities: 300 g – 375 g, 12-15 weeks) and virgin female (body weight and age at arrival in animal facilities: 200 g – 230 g, 12-15 weeks) Sprague-Dawley rats were purchased from Envigo (Indianapolis, IN, USA). After arrival to the animal facilities, rats were acclimatized for one week before any experimentation. All rats were double-housed under 12:12 h light/dark cycle in a temperature and humidity-controlled environment and had free access to water and standard laboratory chow throughout the study. After the period of acclimatization, a normal estrous cycle was determined using vaginal cytology. Females in late proestrus/early estrous were mated with a male overnight (pair mating). The following morning, if spermatozoa were detected in vaginal smears, that day was designated as day 1 of gestation (GD1; term = 22-23 days) (41, 42). Final experiments were performed when rats were 18-24 weeks old. In total, 45 female rats (n=8 non-pregnant, n=37 pregnant) were used. Euthanasia and tissue harvest were performed at 9:00 am and vascular reactivity experiments were performed immediately after.

### Tissue Harvest

Pregnant rats (G16; term = 22-23 days) and age-matched non-pregnant rats were euthanized by isoflurane overdose followed by cutting the diaphragms and removal of their hearts. Uterine horns (including fetuses and placentas for pregnant rats) with uterine arteries and surrounding adipose tissue were excised and placed in ice-cold Krebs-Henseleit solution with the following composition (in mM): 130 NaCl, 4.7 KCl, 14.9 NaHCO_3_, 5.5 dextrose, 1.18 KH_2_PO_4_, 1.17 MgSO_4_·7H_2_O, 1.6 CaCl_2_, and 0.026 EDTA (all obtained from Sigma-Aldrich).

Fetuses were euthanized via decapitation. After carefully removing the surrounding adventitia and PVAT, uterine arterial segments were immediately used for vascular reactivity experiments or functional bioassays. Functional bioassays were used to determine vascular release of TxA_2_ and PGI_2_. PVAT was used to prepare PVAT-conditioned media (PVATmedia) as previously described (41). Briefly, PVATmedia was prepared by incubating 0.4 mg of uterine PVAT in 15 mL Krebs-Henseleit solution, which was continuously gassed with 95% O_2_ and 5% CO_2_. Following 1 hour incubation in Krebs-Henseleit solution, PVATmedia was filtered with a 0.20 µm syringe filter (cat#: 431222, Corning Life Sciences, Corning, NY) and the filtrate was added to the myograph chambers for vascular reactivity experiments used for uterine artery stimulation (41).

### Vascular reactivity experiments

PVAT-denuded uterine arteries were cut in 2 mm rings and mounted onto a multiwire small vessel myograph system (620M, Danish Myo technology A/S, Aarhus, Denmark).

Myograph chambers were filled with Krebs-Henseleit solution (37°C), which was continuously gassed with 95% O_2_ and 5% CO_2_. Optimal baseline tension was determined for each individual vascular segment using the DMT Normalization Module for LabChart Software (ADInstruments, Colorado Springs, CO) as previously described (41, 42). Vessels were allowed to equilibrate for 1 hour at baseline tension. Arteries were twice exposed to potassium chloride (KCl, 120 mM) in order to evaluate vascular integrity. After washing and returning to baseline tension, vessels were exposed to a bolus of PE (1 µM) followed by a bolus of ACh (3 µM) to assess endothelial viability. Vessels with < 80% relaxation to ACh were excluded from the experimental protocol and analysis. Subsequent concentration-response curves were carried out after vessels were washed with Krebs-Henseleit solution and baseline tension was restored.

To determine the effect of PVATmedia on the contribution of COX- and NOS-dependent pathways to ACh-induced relaxation, ACh concentration-response curves (1 nM – 30 µM) were performed in arteries exposed to Krebs-Henseleit solution (control) or PVATmedia in the presence of one of the following inhibitors: a) indomethacin, 10 µM; b) SC560, 1 µM (selective COX-1 inhibitor; IC_50_ = 9 nM for recombinant COX-1, IC_50_ = 6.3 µM for recombinant COX-2 (56); c) NS398, 1 µM (selective COX-2 inhibitor; IC_50_ = 1.77 µM for recombinant COX-2, IC_50_ = 75 µM for recombinant COX-1 (6)); d) L-NAME, (non-selective NOS inhibitor, 100 µM); SQ 29,548 (selective thromboxane receptor (TP) antagonist, K_B_ = 0.5 – 1.7 nM (40)). PVATmedia was added in the myograph chamber for 10 minutes before the inhibitors, which were added in the chamber 30 minutes before the ACh concentration-response curves were generated. None of the arterial segments were incubated with more than one inhibitor in each experiment to avoid carryover effects.

### Release of prostanoids from uterine arteries

Concentrations of thromboxane B_2_ (TxB_2_) and 6-keto Prostaglandin F1α (PGF_1α_), which are stable metabolites of TxA_2_ and PGI_2_, respectively, were measured in media from vascular tissues as previously described (42). Briefly, uterine arteries were incubated in 500 µL of Krebs-Henseleit solution or PVATmedia for 30 min at 37°C. L-NAME (100 µM) was included in all incubations because previous studies have shown that endogenous NO modulates the COX pathway and the release of COX-derived prostanoids (52). At the end of the incubation period, Krebs-Henseleit solution and PVATmedia were collected, snap frozen in liquid nitrogen and stored at −80°C for subsequent measurements of TxB_2_ and 6-keto PGF_1α_ concentrations. TxB_2_ and 6-keto PGF_1α_ concentrations were measured using ELISA as instructed by the manufacturer and as we previously published (42). Samples were used in 1:10 and 1:3364 (two serial dilutions at 1:58 each) dilution for the TxB_2_ and 6-keto PGF_1α_ assays, respectively. The intra- and inter-assay coefficients of variation were smaller than 16%. Sample concentrations are expressed as picograms per ml per mg of tissue wet weight (pg/ml/mg tissue).

### Western blot analysis

Western blot analysis was used to assess protein expression in arteries from two different protocols: a) arteries incubated with PVATmedia or Krebs-Henseleit solution for 30 min, b) arteries from the functional bioassays described above, in which vessels were incubated with Krebs-Henseleit solution plus L-NAME or PVATmedia plus L-NAME.

Uterine arteries were homogenized in ice-cold T-PER lysis buffer containing sodium orthovanadate, 100 mM PMSF and a tablet of protease inhibitor cocktail. Protein concentration was determined with a modified BCA protein assay using the following reagents: Pierce™ Bovine Serum Albumin Standard, Pierce™ BCA Protein Assay Reagent A and Pierce™ BCA Protein Assay Reagent B. Samples containing 7 µg protein were resolved by electrophoresis on 10% SDS-PAGE gels (1 hour at 80 V, 1 hour at 100 V at room temperature) and then transferred to nitrocellulose membranes (overnight, 25 V, 4°C). Membranes were probed for the vascular content of COX-1 (anti-COX-1, cat#: 4841S, source: rabbit, 1:2000; Cell Signaling Technology, Danvers, MA, USA) and COX-2 (anti-COX-2: cat#: 610204, source: mouse, 1:1000; BD Biosciences, San Jose, CA, USA). The immunostaining was detected using horseradish peroxidase-conjugated anti-rabbit immunoglobulin G (IgG) (GE Healthcare, Buckinghamshire, UK; dilution: 1:5000) or anti-mouse IgG (GE Healthcare; 1:5000) for 1 hour at room temperature. Results were normalized to the expression of β-actin (cat#: A3854, source: mouse, 1:80,000, Sigma-Aldrich; anti-mouse IgG (GE Healthcare; dilution: 1:25,000). Immunoreactive bands were visualized with an enhanced chemiluminescence detection system (ChemiDoc Imaging System, BIO-RAD, Hercules, CA) and quantified using Image Lab Software (BIO-RAD). The original uncropped western blots are shown in supplemental materials.

## Data analysis

### Concentration-response curves

Concentration-response curves were fit and analyzed using robust nonlinear regression analysis as previously described (GraphPad Prism software, v. 8.4, San Diego, CA, USA) (41). EC_50_ (expressed as *p*EC_50_: negative logarithm of EC_50_) and Emax were calculated for each sigmoidal curve. Area under the curve (i.e., cumulative relaxation response; AUC) was used to evaluate the modulatory effect of PVATmedia and inhibitors on vascular response profiles. Relaxation responses are expressed as a percentage of maximum response to PE.

### Statistical analysis

Concentration-response curves are presented as mean ± SEM. *p*EC_50_, Emax, and AUC data are presented in scatter plots with means, SEM, and individual data points. *p*EC_50_, Emax, AUC were analyzed using one-way analysis of variance (ANOVA) followed by a Tukey post hoc test. Paired t-tests were used to compare concentrations of TxB_2_ and 6-keto PGF_1α_ and COX-1 and COX-2 protein expression between uterine arteries (from the same animal) incubated with and without PVATmedia. All statistical analyses were conducted with Prism software (v. 8.4). The significance level was set at *α* = 0.05 for all comparisons.

## Results

Relaxation responses to ACh were lower in uterine arteries incubated with PVATmedia [(+)PVATmedia] compared to control arteries [(-)PVATmedia] in pregnant rats (Fig. 1A). Indomethacin reduced relaxation responses to ACh in control arteries (Fig. 1B), indicating that COX-derived relaxing factors contributed to ACh-induced relaxation in isolated uterine arteries. Contrary, in arteries incubated with PVATmedia, indomethacin augmented relaxation responses to ACh (Fig. 1C), suggesting the involvement of COX-derived constricting factors in the anti-dilatory effects of PVATmedia. E_max_ (Fig. 1D), *p*EC_50_ (Fig. 1E), and AUC (Fig. 1F) remained lower in arteries incubated with PVATmedia plus indomethacin as compared to control arteries (i.e., no PVATmedia, no indomethacin), indicating the contribution of a non-COX-related mechanism to the anti-dilatory effects of PVATmedia. The effects of PVATmedia and indomethacin found in arteries from pregnant rats were not observed in uterine arteries from non-pregnant rats, where neither PVATmedia nor indomethacin alone reduced relaxation responses to ACh (Fig. 2A-F). L-NAME abolished ACh-induced relaxation in control uterine arteries from pregnant rats (Fig. 3A-B), and this effect remained significant in the presence of PVATmedia (Fig. 3C-E). These data suggested PVATmedia affected COX-but not NOS-dependent contributions to ACh-induced relaxation in isolated uterine arteries from pregnant rats.

**Figure 1.**
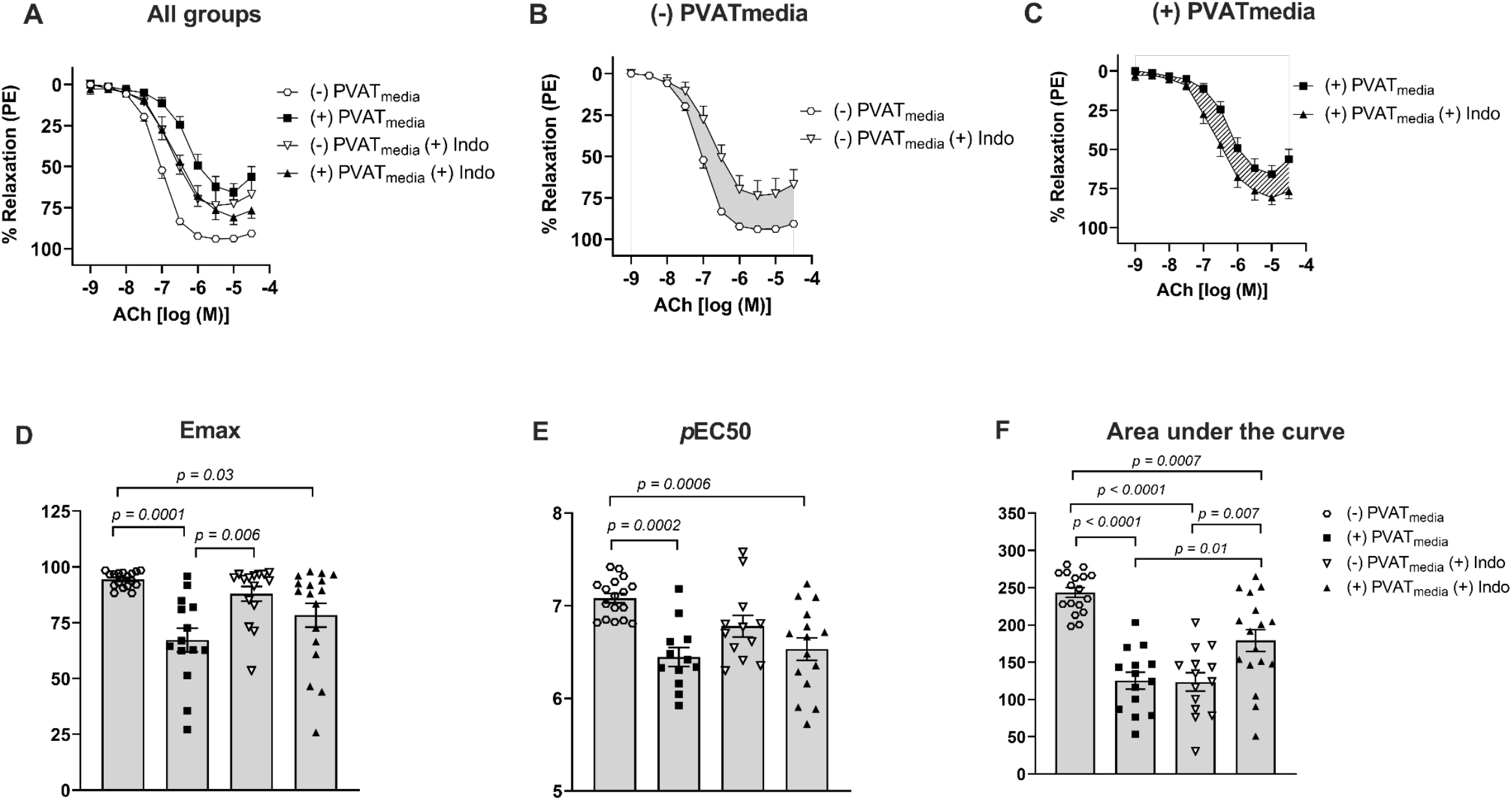
Effects of COX inhibition on ACh-induced relaxation in uterine arteries exposed to PVAT-derived factors in pregnant rats. (A) All treatments; (B) Concentration-response curves to ACh in the presence or absence of indomethacin (no PVATmedia added). Grey shaded area indicates a reduction in ACh-induced relaxation in response to indomethacin; (C) Concentration-response curves to ACh in the presence or absence of indomethacin (PVATmedia added). Pattern-filled area indicates an increase in ACh-induced relaxation in response to indomethacin; (D) Maximum responses (Emax) to ACh for all concentration-response curves; (E) Negative logarithm of EC_50_ (*p*EC_50_) for all concentration-response curves; (F) Area under the curve for all concentration-response curves. Values are mean ± SEM. One-way ANOVA followed by Tukey’s post hoc test. ACh, acetylcholine; COX, cyclooxygenase; PVATmedia, perivascular adipose tissue conditioned media.

**Figure 2.**
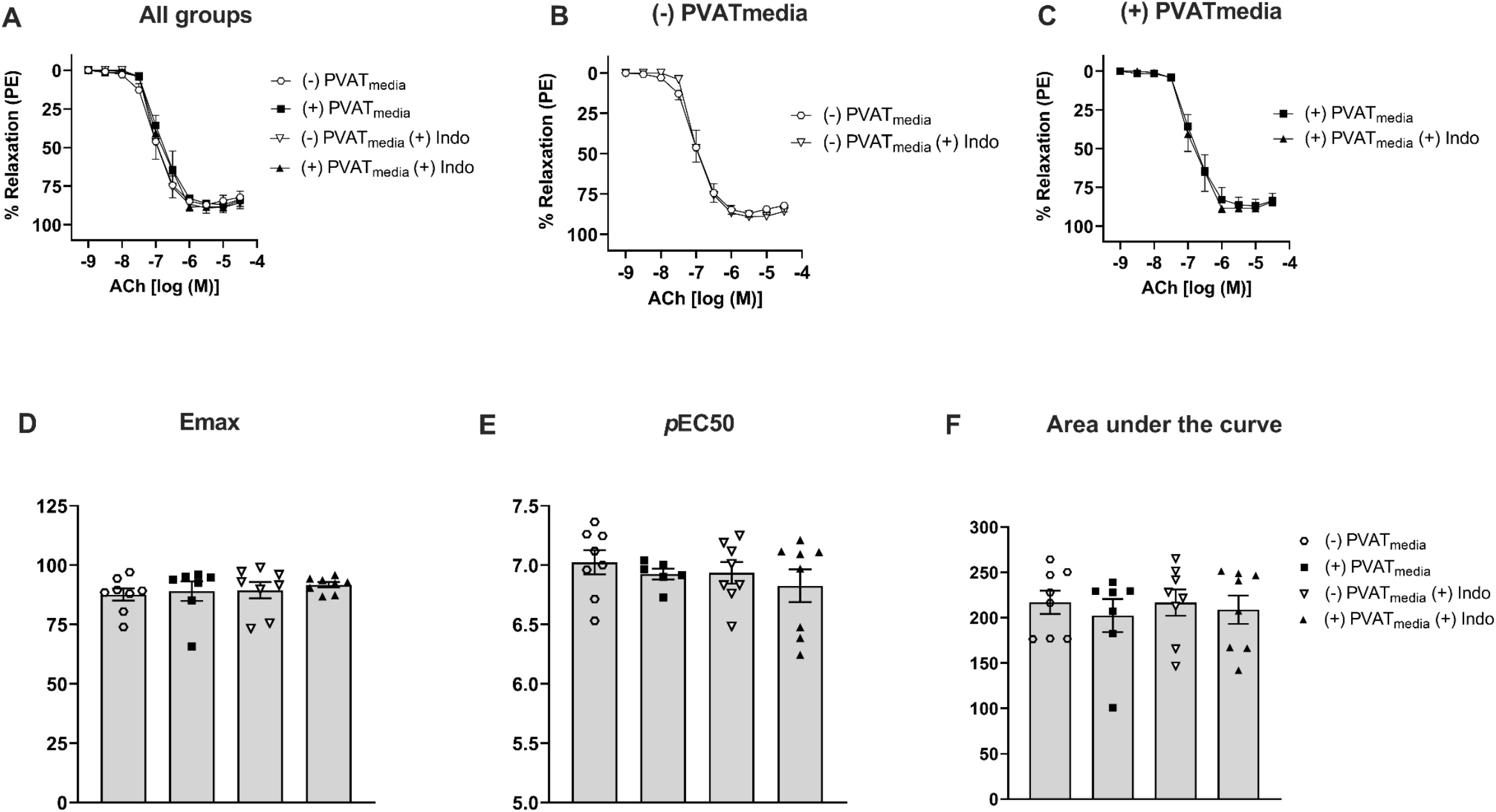
Effects of COX inhibition on ACh-induced relaxation in uterine arteries exposed to PVAT-derived factors in non-pregnant rats. (A) All treatments; (B) Concentration-response curves to ACh in the presence or absence of indomethacin (no PVATmedia added); (C) Concentration-response curves to ACh in the presence or absence of indomethacin (PVATmedia added); (D) Maximum responses (Emax) to ACh for all concentration-response curves; (E) Negative logarithm of EC_50_ (*p*EC_50_) for all concentration-response curves; (F) Area under the curve for all concentration-response curves. Values are mean ± SEM. One-way ANOVA followed by Tukey’s post hoc test. ACh, acetylcholine; COX, cyclooxygenase; PVATmedia, perivascular adipose tissue conditioned media.

**Figure 3.**
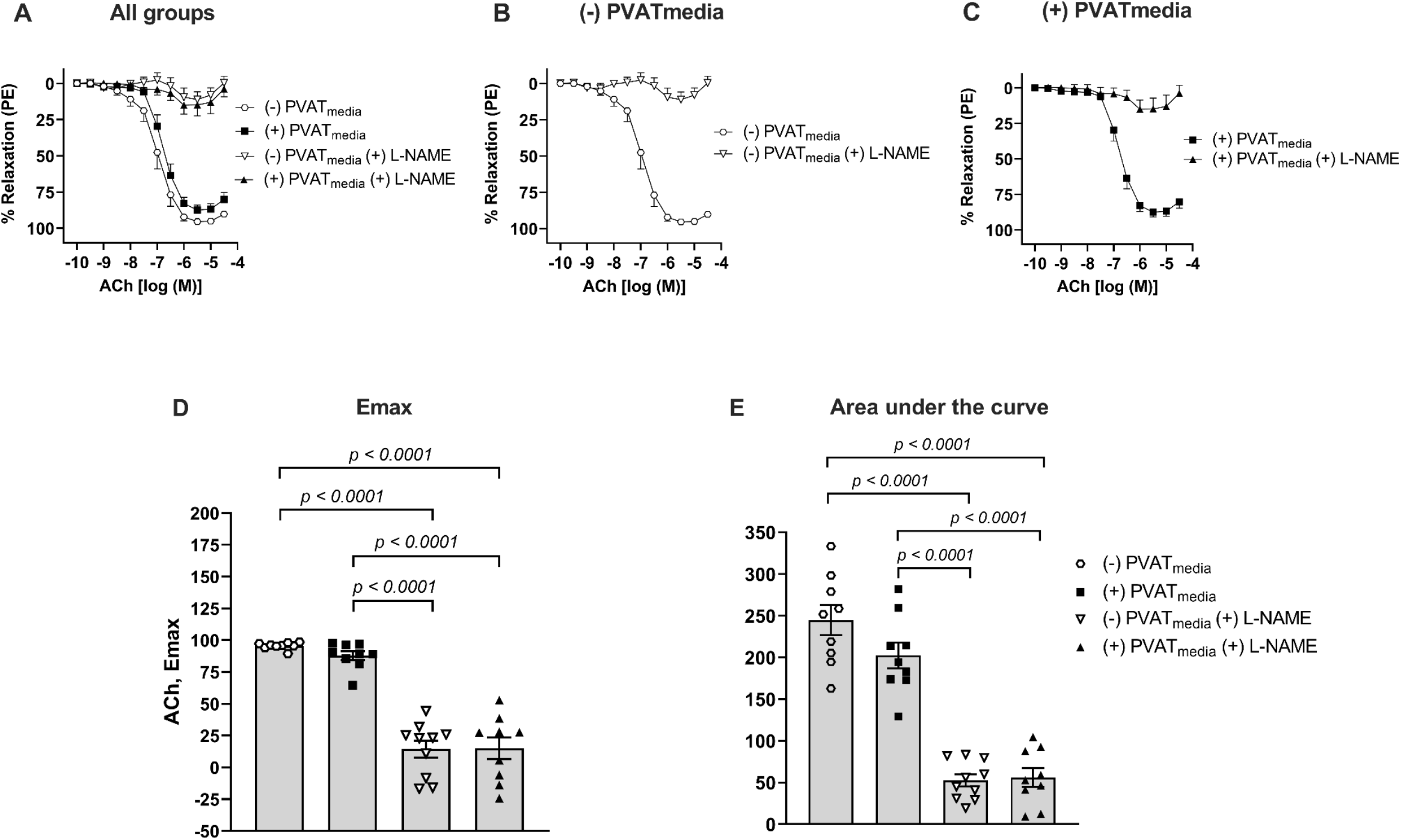
Effects of NOS inhibition on ACh-induced relaxation in uterine arteries exposed to PVAT-derived factors in pregnant rats. (A) All treatments; (B) Concentration-response curves to ACh in the presence or absence of L-NAME (no PVATmedia added); (C) Concentration-response curves to ACh in the presence or absence of L-NAME (PVATmedia added); (D) Maximum responses (Emax) to ACh; (E) Area under the curve for all concentration-response curves. Values are mean ± SEM. One-way ANOVA followed by Tukey’s post hoc test. ACh, acetylcholine; NOS, nitric oxide synthase; PVATmedia, perivascular adipose tissue conditioned media.

To determine which COX isoform is involved in the anti-dilatory effect of uterine PVAT, we incubated uterine arteries with SC560 (COX-1 inhibitor; Fig. 4A-C) or NS398 (COX-2 inhibitor; Fig. 4D-F) in the presence or absence of PVATmedia. Since the effects of indomethacin on the anti-dilatory influences of PVATmedia were not observed only in non-pregnant rats, these experiments concentrated on uterine arteries from pregnant rats. SC560 (Fig. 4A, C) reduced relaxation to ACh in naive uterine arteries. In contrast, NS398 did not affect total relaxation (AUC) to ACh in these vessels [(-)PVATmedia vs. (-)PVATmedia(+)NS398, p=0.61]. These data indicated that COX-1 but not COX-2-derived products contributed to ACh-induced relaxation in isolated control uterine arteries. In contrast, in uterine arteries incubated with PVATmedia, neither SC560 (Fig. 4B, C) nor NS398 (Fig. 4E, F) affected relaxation responses to ACh [AUC: (+)PVATmedia vs. (+)PVATmedia(+)SC560, p=0.85; (+)PVATmedia vs. (+)PVATmedia(+)NS398, p=0.99].

**Figure 4.**
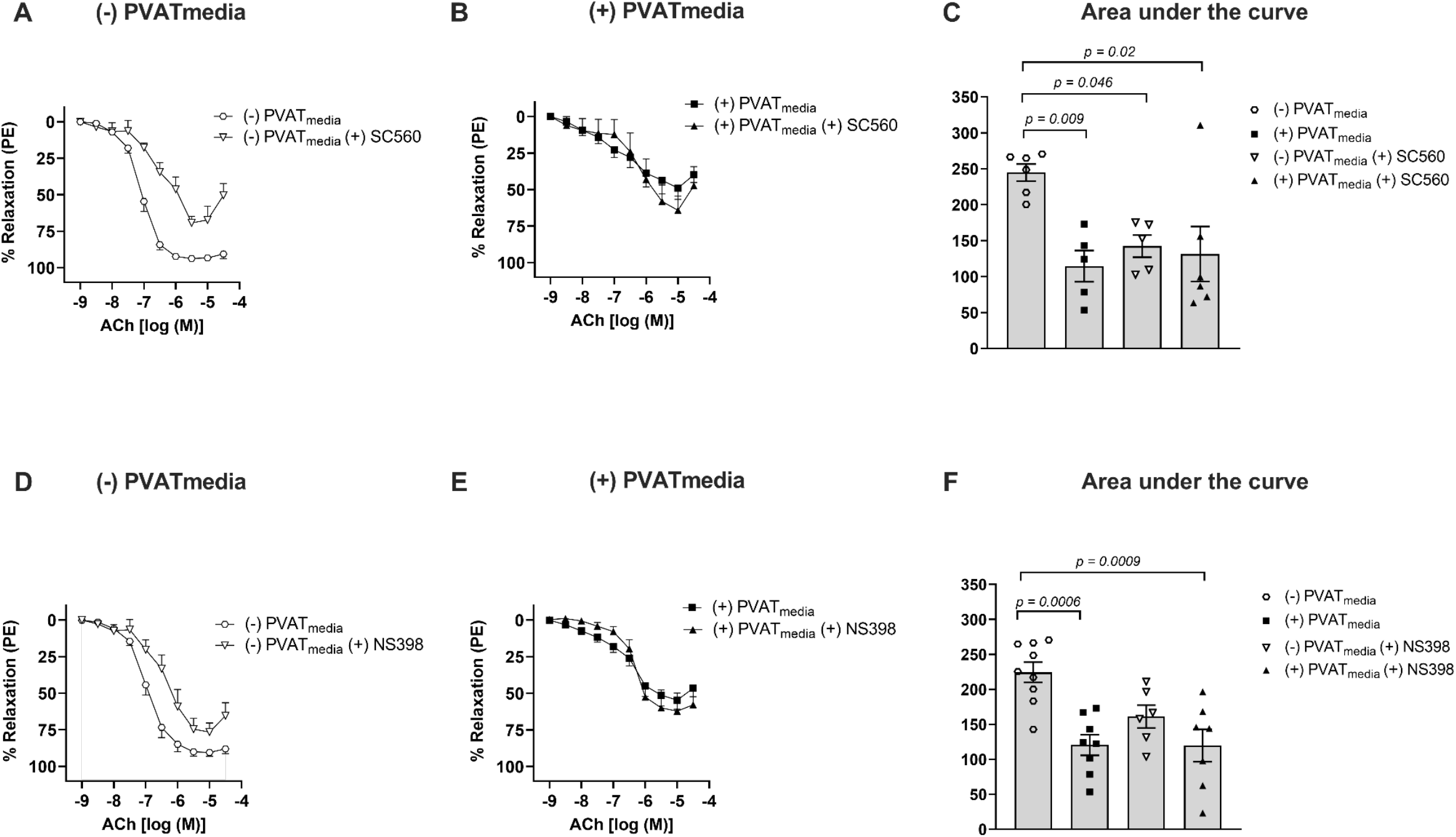
Effects of inhibition of COX isoforms, COX-1 and COX-2, on ACh-induced relaxation in uterine arteries exposed to PVAT-derived factors in pregnant rats. (A) Concentration-response curves to ACh in the presence or absence of COX-1 inhibitor, SC560 (no PVATmedia added); (B) Concentration-response curves to ACh in the presence or absence of SC560 (PVATmedia added); (C) Area under the curve for all concentration-response curves in (A) and (B); (D) Concentration-response curves to ACh in the presence or absence of COX-2 inhibitor, NS398 (no PVATmedia added); (E) Concentration-response curves to ACh in the presence or absence of NS398 (PVATmedia added); (F) Area under the curve for all concentration-response curves in (D) and (E). Values are mean ± SEM. One-way ANOVA followed by Tukey’s post hoc test. ACh, acetylcholine; COX, cyclooxygenase; PVATmedia, perivascular adipose tissue conditioned media.

Uterine arteries incubated with PVATmedia released greater concentrations of TxB_2_ compared to control arteries (i.e., arteries incubated with Krebs-Henseleit) (Fig. 5A). However, incubation with PVATmedia did not affect 6-keto PGF_1α_ release (Fig. 5C). In this protocol, L-NAME was added in our experiments to control for the potential effects of basal levels of NO on the COX pathway. To determine if an increase in PVAT-mediated TxB_2_ production could explain the anti-dilatory effects of PVAT on uterine arteries from pregnant rats, we performed ACh concentration-response curves in the presence of a TP antagonist. Antagonism of TP with SQ 29,548 did not reverse the anti-dilatory effects of PVATmedia in uterine arteries (Fig. 6A-F).

**Figure 5:**
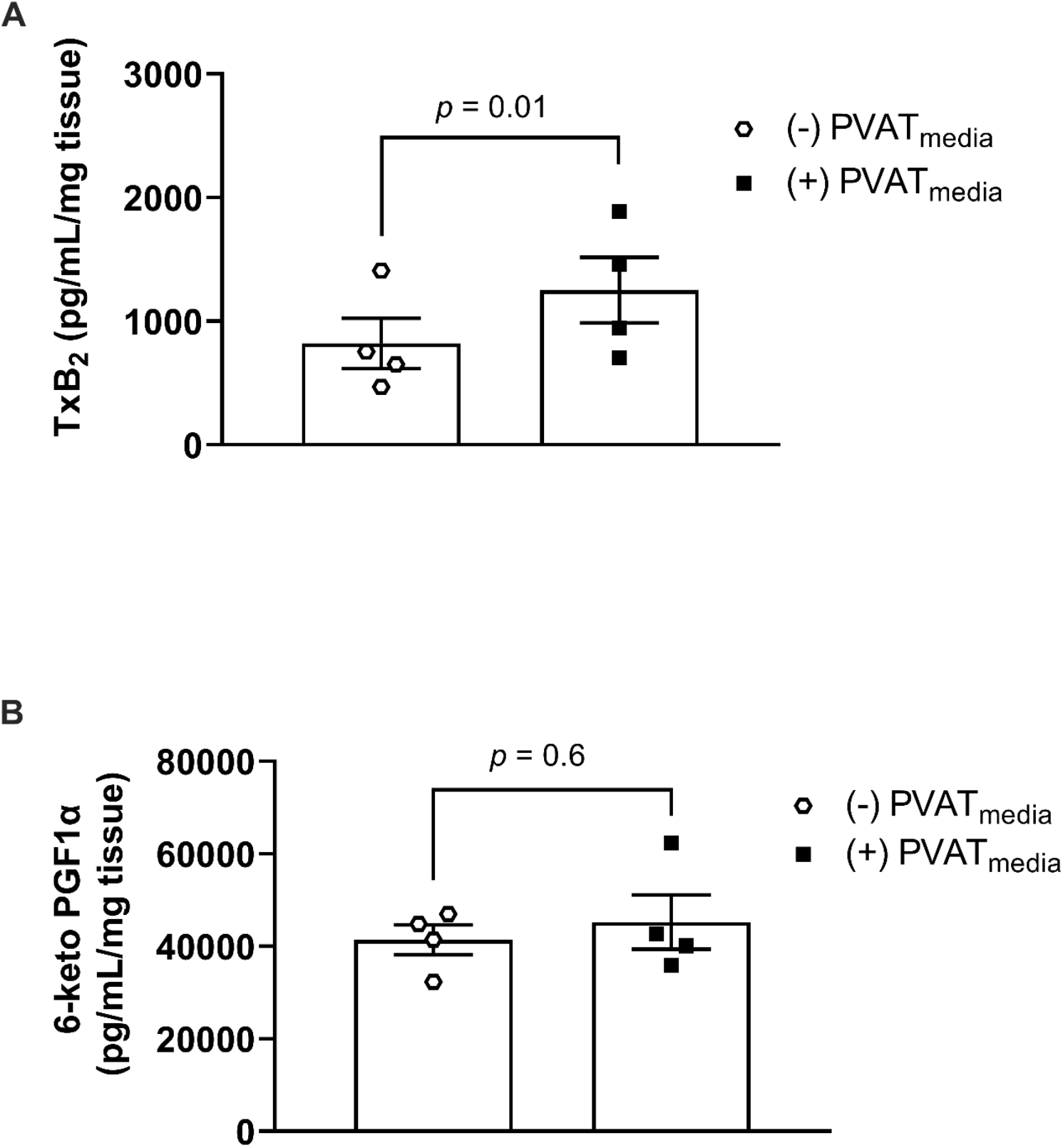
Effects of PVAT-derived factors on prostanoid production. Concentrations of (A) TxB_2_ and (B) 6-keto PGF_1α_ in uterine arteries from pregnant rats. Arteries were incubated in Krebs-Henseleit solution [(-) PVATmedia] or PVATmedia [(+) PVATmedia] in the presence of a nitric oxide synthase inhibitor (L-NAME). Values are mean ± SEM. Paired t-test. COX, cyclooxygenase; PVATmedia, PVATmedia, perivascular adipose tissue conditioned media; TxB_2_, thromboxane B_2_; 6-keto PGF_1α_, 6-keto prostaglandin F1α.

**Figure 6.**
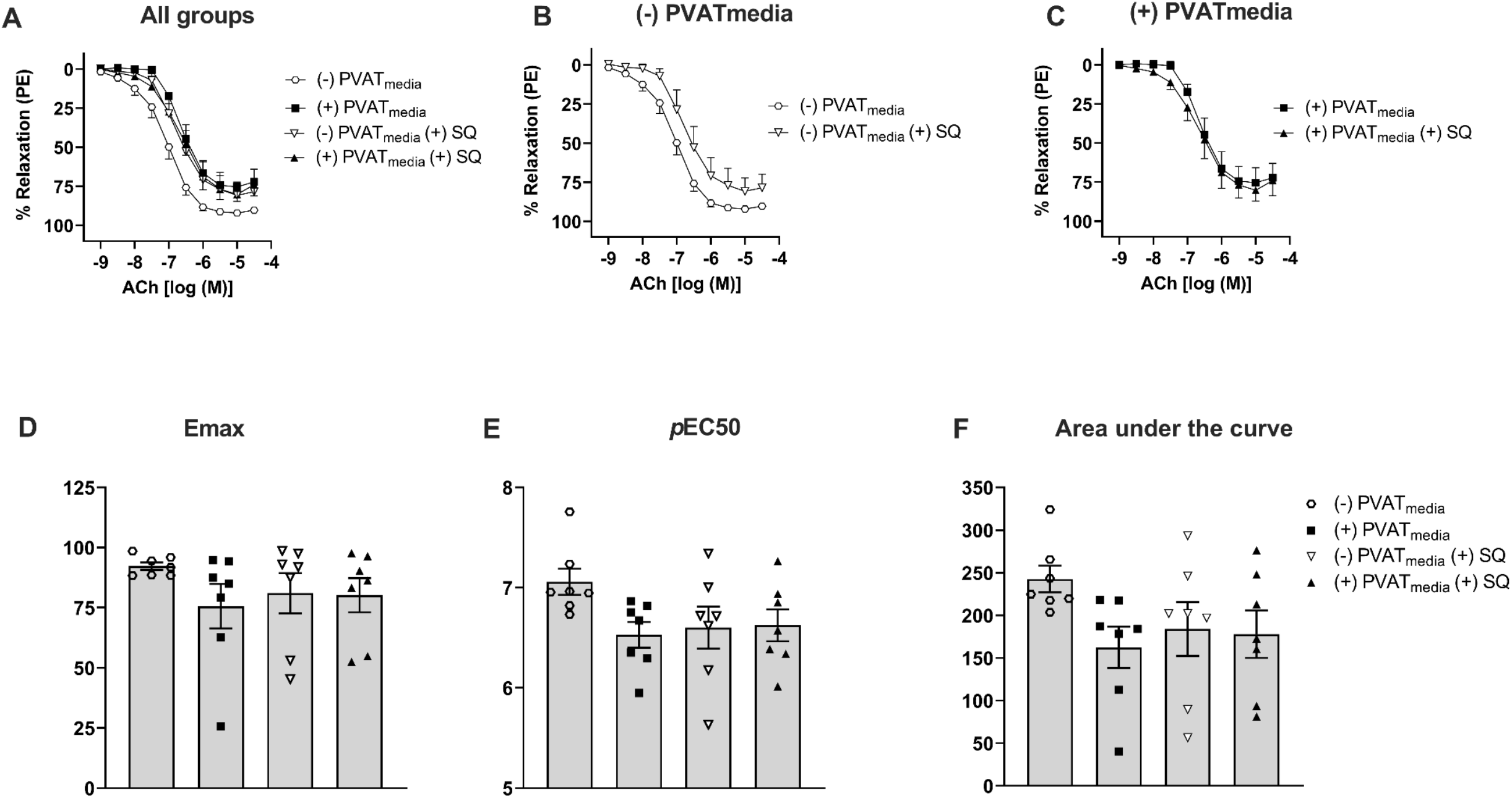
Effects of TP receptor inhibition on ACh-induced relaxation in uterine arteries exposed to PVAT-derived factors in pregnant rats. (A) All treatments; (B) Concentration-response curves to ACh in the presence or absence of SQ 29,548 (no PVATmedia added); (C) Concentration-response curves to ACh in the presence or absence of SQ 29,548 (PVATmedia added); ((D) Maximum responses (Emax) to ACh for all concentration-response curves; (E) Negative logarithm of EC_50_ (*p*EC_50_) for all concentration-response curves; (F) Area under the curve for all concentration-response curves. Values are mean ± SEM. One-way ANOVA followed by Tukey’s post hoc test. ACh, acetylcholine; PVATmedia, perivascular adipose tissue conditioned media; TP, thromboxane receptor.

Exposure to uterine PVATmedia increased COX-1 and COX-2 protein expression (Fig. 7A-B, Fig. S1) in uterine arteries pre-treated with L-NAME, while it had no effect on COX-1 and COX-2 protein expression in arteries that were not treated with L-NAME (Fig. S2-3).

**Figure 7:**
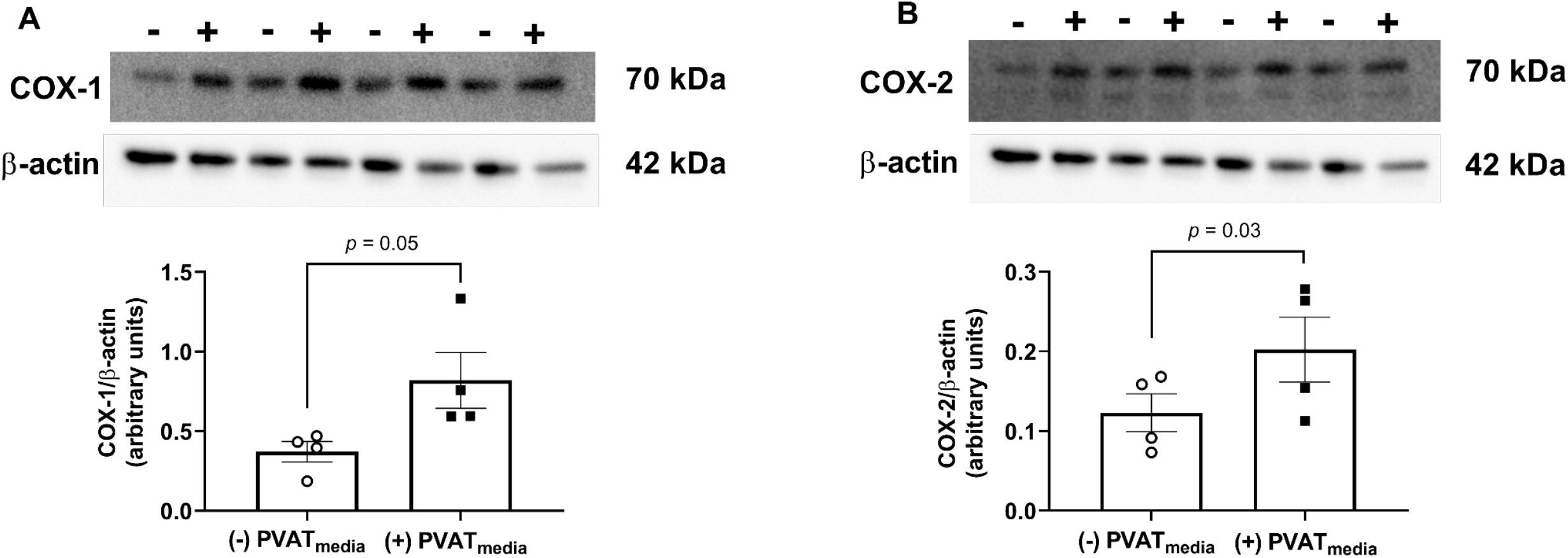
Effects of PVAT-derived factors on COX protein content. Relative protein expression of (A) COX-1 and (B) COX-2 in uterine arteries from pregnant rats. Arteries were incubated in Krebs-Henseleit solution [(-) PVATmedia)] or PVATmedia [(+) PVATmedia] in the presence of a nitric oxide synthase inhibitor (L-NAME). Values are mean ± SEM. Paired t-test. COX, cyclooxygenase; PVATmedia, PVATmedia, perivascular adipose tissue conditioned media; TxB_2_, thromboxane B_2_; 6-keto PGF_1α_, 6-keto prostaglandin F1α.

## Discussion

The main finding of this study was that inhibition of the COX pathway suppressed the anti-dilatory effects of PVATmedia on uterine arteries from pregnant rats, while PVATmedia had no effect on the contribution of the NOS/NO pathway to ACh-induced relaxation in these vessels. Our data suggest that PVAT releases factors that may potentiate the production of COX-derived constrictor products, which oppose the dilatory effects of ACh in uterine arteries. In addition, we found that a COX-mediated pathway contributes to ACh-induced relaxation in isolated uterine arteries from pregnant but not non-pregnant rats, indicating that pregnancy may upregulate COX signaling in this vascular preparation.

In most vascular beds, PVAT has anti-contractile effects (5, 23). In isolated swine coronary arteries, however, PVAT is pro-contractile (39, 44). Recently, we reported that PVAT was also pro-contractile in uterine arteries from healthy pregnant rats (41), confirming the notion that the vasoactive effects of PVAT depend on the vascular region and physiological state (5, 23). In addition, PVAT had anti-dilatory actions in maternal uterine arteries as it suppressed relaxation responses to ACh (41). We attributed the vasoactive effects of uterine PVAT to its paracrine actions, since conditioned media prepared from PVAT had pro-contractile and anti-dilatory effects (41). Others have shown that the vasoactive properties of PVAT are due to its ability to uptake neurotransmitters or the ability of its adipocytes to release transferable molecules with vasoactive properties (i.e., PVAT-derived factors) (15, 41, 49). In the present study, we replicated our previous findings (41) as we demonstrated that ACh-induced relaxation was reduced in the presence of PVAT-derived factors. The anti-dilatory effects of PVAT during pregnancy are unique to the reproductive vascular bed as PVAT in mesenteric arteries from pregnant rats reduced contractile responses of isolated pressurized mesenteric arteries (35), while aortic PVAT had anti-contractile effects that were abolished during rat pregnancy (1, 2).

Previously, we observed the endothelium was required for the anti-dilatory effects of PVAT in uterine arteries from pregnant rats, and this observation led to the current investigation. Another observation was that uterine PVAT from pregnant rats lost its vasoactive function when incubated with uterine arteries from non-pregnant rats, while uterine arteries from pregnant rats remained responsive to PVAT from their non-pregnant counterparts.

Together, these observations suggest: a) PVAT-derived factors affect the contribution of the endothelium to vascular smooth muscle relaxation and b) the anti-dilatory effects of PVAT are mediated, at least in part, by pregnancy-induced adaptations of the maternal uterine vascular wall. Thus, we sought to understand whether PVAT-derived factors impact the contribution of endothelium-mediated vasodilatory components of ACh-induced relaxation in uterine arteries.

Previous studies have shown that PVAT suppresses ACh-induced relaxation in canine coronary arteries by inhibiting eNOS (47). Specifically, Payne et al. demonstrated that coronary PVAT releases leptin, which stimulates protein kinase-C beta (PKC-β) phosphorylation of eNOS at its inhibitory site, Thr495, thus inhibiting eNOS, reducing NO production and diminishing coronary artery relaxation (47, 48). Others have shown that PVAT increased Caveolin-1 protein expression (30), which could lead to inhibition of eNOS and reduction of NO generation (30). The eNOS/NO pathway is the predominant contributor of relaxation responses to ACh in large vessels (55); thus, we hypothesized that PVAT may release factors that inhibit this pathway and contribute to reduced ACh-induced relaxation. In contrast to previous investigations of non-reproductive vascular beds, we demonstrated PVAT-derived factors did not affect the contribution of eNOS/NO pathway to ACh-induced relaxation in uterine arteries from pregnant rats. In addition, when we inhibited the COX pathway in the presence of an intact and unopposed eNOS system, the relaxation response to ACh was increased by PVAT. These data further demonstrate that PVAT has no inhibitory effect on the eNOS/NO pathway in uterine arteries from pregnant rats. Previously we found discrepancies between outcomes from in vitro compared to in vivo experiments showing that PVAT potentiated UtBF during rat pregnancy, while it reduced ACh-induced relaxation in isolated rat uterine arteries. We suggested that what we observed was the individual effect of PVAT on the uterine vascular wall in isolated arteries, while the *in vivo* experiments addressed the net effect of PVAT, which may be modulated by systemic and hemodynamic responses. Although delineating the differences between the *in vitro* and *in vivo* studies is above and beyond the scope of the present investigation, these differences should be taken into account when interpreting the pharmacological manipulation of isolated vascular tissues, especially in the context of the NOS/NO and COX pathways, which are greatly affected by local shear stress.

Our findings in control uterine arteries revealed that COX-derived products have a significant contribution to relaxation responses to ACh in uterine arteries. In addition, we found that this dependency on COX-derived factors was specific to pregnancy because COX inhibition had no effect in uterine arteries from non-pregnant rats, suggesting pregnancy-induced upregulation of production of COX-derived relaxing factors in response to ACh stimulation. As a result of this observation, we focused on examining the effect of PVAT on the contribution of COX pathway to ACh-induced relaxation. When we investigated the effect of PVAT on COX-mediated ACh-induced relaxation, we found that COX inhibition increased relaxation responses to ACh in arteries exposed to PVATmedia. This is the opposite effect of what we found in control uterine arteries, leading us to conclude that PVAT-derived factors may inhibit the production of COX-derived relaxing factors or it may potentiate the production of COX-derived constrictor products in the maternal uterine vascular wall from pregnant rats.

To the best of our knowledge, this is the first study to a) demonstrate the importance of the COX pathway in ACh-induced relaxation in isolated uterine arteries in rat pregnancy and b) report the anti-dilatory effects of uterine PVAT-derived factors are mediated, at least in part, by COX-derived products.

COX enzymes are key mediators of the conversion of arachidonic acid to vasoactive prostanoids, such as PGI_2_ and TxA_2_ (34). The production of PGI_2_ increased 2-3 fold in uterine arteries and uterine artery endothelial cells from pregnant ewes (24, 28, 33). This increase was attributed to upregulation of COX-1 in response to increased pregnancy-associated hemodynamic and hormonal changes (34). There are two main isoforms of COX enzymes (16), COX-1 and COX-2, both of which are inhibited by indomethacin (58). We further investigated the contribution of each enzyme to the anti-dilatory effects of PVAT using selective pharmacological inhibition. Inhibition of COX-1 reduced relaxation responses to ACh in control uterine arteries from pregnant rats, whereas inhibition of COX-2 had no effect. This finding suggests that in isolated uterine arteries from pregnant rats, ACh-induced relaxation is in part depended on COX-1-derived products, while the COX-2 pathway did not have a significant contribution to this response.

Selective inhibition of COX-1 or COX-2 did not prevent the anti-dilatory effects of PVAT, even though non-selective COX inhibition with indomethacin reversed these effects. There are several explanations for the discrepancy between selective and non-selective pharmacological inhibition. Previous studies showed that indomethacin increased ACh-induced relaxation in rat aortic rings via a COX-independent that involved superoxide anion generation (13). In addition, the contribution of another COX-dependent mechanism to this anti-dilatory effect may also be present. COX-3 is the most recently identified COX isoform (9). COX-3 is a COX-1 variant encoded by the *PTGS1* gene and shares catalytic and structural features of COX-1 (9). The IC_50_ value of indomethacin is similar for COX-1 (0.01 µM) and COX-3 (0.016 µM) and greater for COX-2 (0.66 µM) (9), while the effects of SC560 and NS398 on COX-3 are unclear. Erol et al. demonstrated that inhibition of COX-3 increased dilatory responses to bradykinin in rat aorta (14), suggesting the contribution of COX-3 to vasodilation in this vascular bed. In addition to COX-3, small COX-1 derived proteins (PCOX-1) have also been identified in the canine cerebral cortex (9). Whether COX-3 or PCOX-1 are involved in uterine artery responses to ACh and if pregnancy affects their expression and activity are currently unknown.

In these studies, we found that PVAT-derived factors increased the production of TxB_2_ but did not affect the production of PGI_2_ metabolites. These results suggest that PVAT-derived factors may potentiate the generation of COX-derived constrictor products, which oppose the dilatory effects of ACh in uterine arteries. It is noteworthy that there is an interaction between NOS- and COX-dependent pathways (22, 34, 50, 62). To avoid the crosstalk between these two pathways in response to exposure to PVAT-derived factors, our experiments on the production of TxB_2_ and 6-keto PGF_1α_ were performed in the presence of a NOS inhibitor as previously described by us and others (36);(42)}. Interestingly, co-treatment of uterine arteries with PVAT-derived factors and a NOS inhibitor caused an increase in the protein content of COX-1 and COX-2, while PVAT-derived factors alone did not have the same effect. It is possible that the factor(s) secreted by PVAT act as a substrate of vascular COX, promoting prostanoid production, and that NO modulates this property.

TxB_2_ generated in endothelial cells (51) potentiates vascular contraction upon binding to its receptor on vascular smooth muscle cells (11). Since uterine arteries exposed to PVATmedia produced greater concentrations of TxB_2_, we pursued to investigate whether the anti-dilatory effects of PVATmedia on uterine arteries could be explained by activation of the COX-1/TxA_2_/TP pathway, which could oppose the dilatory actions of ACh. Our results indicate that although PVAT increased the production of TxB_2_, a by-product of TxA_2_, the anti-dilatory effects of PVAT were not mediated by TP activation. Thus, we conclude that uterine PVAT releases transferable factors, which reduce ACh-induced relaxation in uterine arteries from pregnant rats via a COX-dependent but TP-independent mechanism.

There are a few limitations in the present studies. We observed an increase in COX enzymes in uterine arteries and a role for COX in our experiments; however, neither of our preparations distinguished between the individual layers of the uterine vascular wall and their contribution to these findings. In addition, our studies did not distinguish whether concentrations of 6-keto PGF_1α_ and TxB_2_ reflected prostanoid release from the uterine PVAT itself (10, 45), uterine vascular wall alone, or both. Further, we only measured PGI_2_ and TxA_2_ metabolites, and it is possible that other prostanoids are released in response to COX activation by uterine PVAT-derived factors. Activation of COXs has been also implicated in the production reactive oxygen species (ROS) (46, 59, 63)} and ROS have been shown to potentiate vascular contraction (12, 54, 60). Thus, we cannot discount a potential role of ROS in the anti-dilatory effects of PVAT. We attempted to perform all experiments related to a specific pathway in vessels from the same animals to increase internal validity and aid the interpretation of our studies. Due to the small amount of protein extracted from homogenized rat uterine arteries, we were unable to perform further investigation on the eNOS/NO pathway. Therefore, our studies provided evidence that PVAT does not affect the contribution of NOS/NO to ACh-induced relaxation but did not assess the effects of PVAT on eNOS protein content and activation. These studies concentrated on the NOS and COX pathways, while the effects of PVAT-derived factors on EDH were not assessed. We concentrated on these two pathways because the contribution of EDH to ACh-induced relaxation has been shown to be important in small resistance vessels and not in large arteries such as the main uterine artery. Nevertheless, the effect of uterine PVAT on EDH in this vessel remains unknown. Uterine artery remodeling in rodents occurs between gestational days 15 - 18 (duration of rat pregnancy=22-23 days) (8, 37, 43)}. In addition, our published studies on the vasoactive effects of PVAT and on the effects of pregnancy on adipose tissue protein/gene profile have been performed on GD16 (41). Thus, in the present study, we concentrated on uterine arteries from pregnant rats on GD16 and did not interrogate the temporal adaptations of the PVAT/uterine artery axis throughout gestation.

## Conclusions and Perspectives

This is the first study to demonstrate that indomethacin reverses the anti-dilatory effects of uterine PVAT in uterine arteries from healthy pregnant rats. The findings of our study may provide the basis for future experiments examining the role of uterine PVAT in pregnancies with dysfunctional uterine artery reactivity, impaired NO pathway, and imbalance in production of constrictor and dilatory prostanoids, such as preeclampsia. These studies may be especially pertinent to pregnancies, in which aspirin is indicated as a preventative approach for risk reduction, such as those pregnancies at high risk of developing preeclampsia.

## Acknowledgments

The authors would like to thank Dr. Stephane Bourque (University of Alberta) for reading the manuscript and providing feedback and Dr. Camilla Wenceslau (The University of Toledo) and Dr. Cameron McCarthy (The University of Toledo) for reviewing the graphical abstract (Fig. 8).

**Figure 8.**
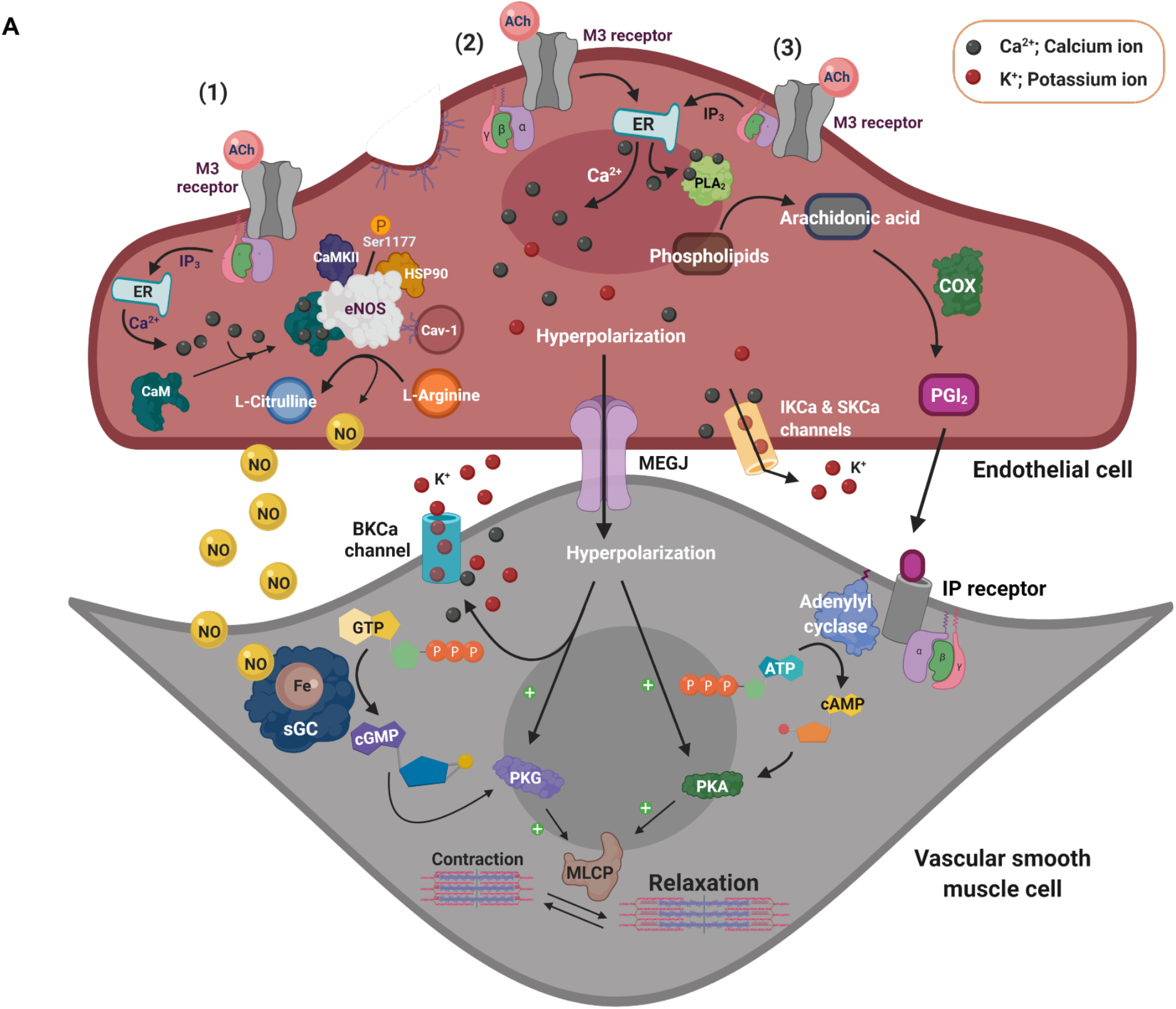
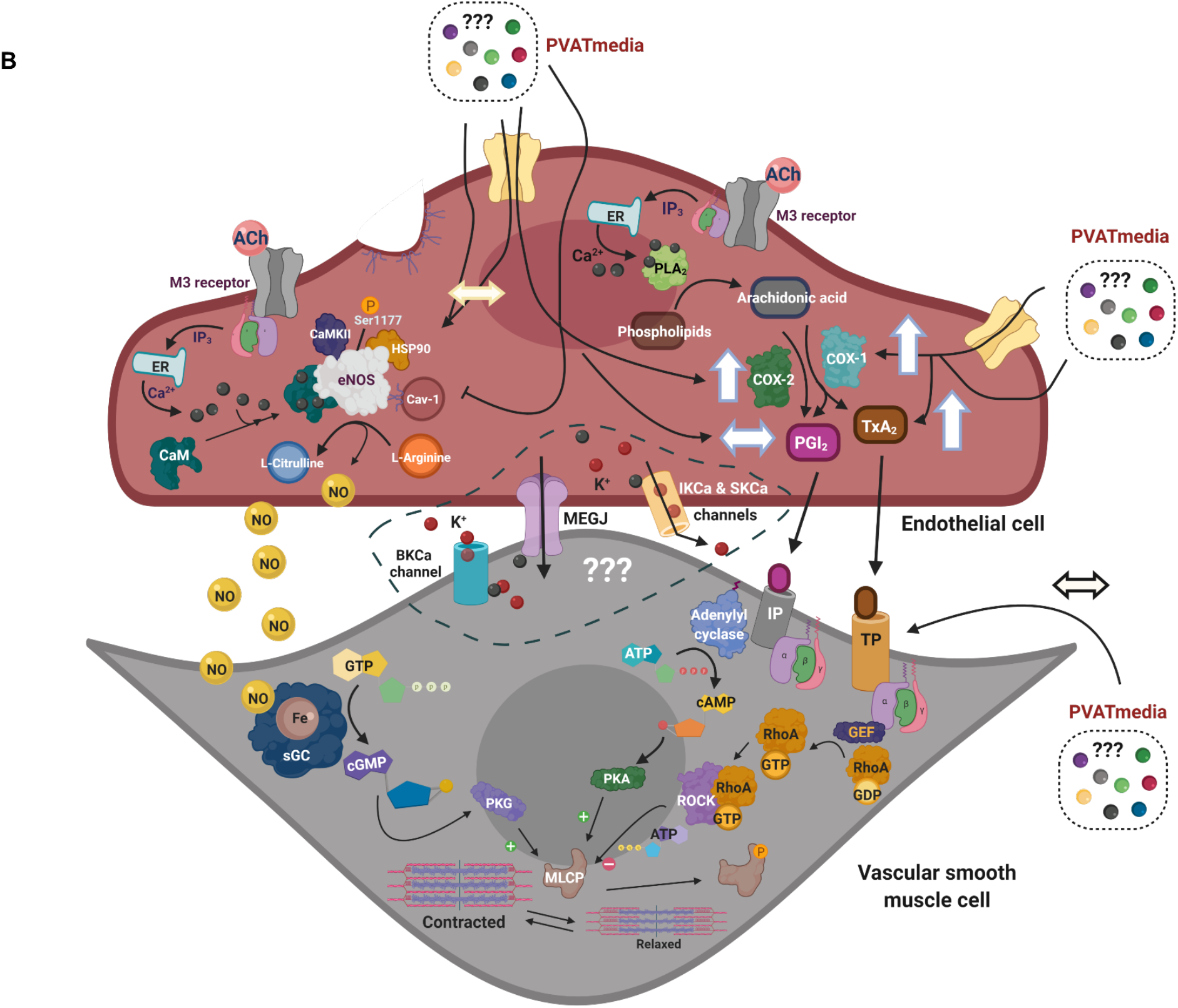
Proposed hypothesis. (A) ACh-induced stimulation of M3 receptors on the vascular endothelium of isolated main uterine arteries induces relaxation of vascular smooth muscle cells via activation of Ca^2+^ dependent mechanisms: (1) eNOS/NO/sGC/cGMP pathway (prominent), (2) EDH pathway, and (3) PLA_2_/COX/PGI_2_/cAMP pathway. In this experimental preparation, uterine arteries are PVAT-denuded and are not exposed to PVAT-derived factors. During pregnancy, there is upregulation of the eNOS/NO pathway and increases in production of COX-derived PGI_2_. (B) Uterine PVAT releases transferable factors that potentiate the COX-1/TxA_2_ pathway shifting the balance between vascular production of TxA_2_ and PGI_2_. These effects may oppose the eNOS/NO mechanism, leading to reduced relaxation responses to ACh in the presence of PVAT-derived factors. M_3_ receptor, Muscarinic acetylcholine receptor 3; ACh, Acetylcholine; ATP, Adenosine-5-triphosphate; ADP, Adenosine-5-diphosphate; BK_Ca_, large-conductance calcium-activated potassium channels; Ca, Calcium; CaM, Calmodulin; cAMP - cyclic Adenosine-5’-monophosphate; CaMKII, Calmodulin kinase II; Cav-1, Caveolin-1; cGMP, cyclic Guanosine-5’-monophosphate; EDH, endothelium-derived hyperpolarization; eNOS, endothelium nitric oxide synthase; Fe, Iron; GDP, Guanosine-5’-diphosphate; GEF, Guanine nucleotide exchange factor; GTP, Guanosine-5’-triphosphate; HSP90, Heat shock protein 90; IP_3_R, IP3 receptor; K+, Potassium; MLCP, Myosin light-chain phosphatase; NO, nitric oxide; PLA_2_, Phospholipase A2; COX, Cyclooxygenase; PGI_2_, Prostaglandin I2; RhoA, Ras homolog family member A; ROCK, Rho-associated protein kinase; sGC, soluble guanylyl cyclase; PKG, Protein kinase G; PKA, Protein kinase A; IP, Prostaglandin I2 receptor; IK, intermediate-conductance calcium-activated potassium channels; SERCA, sarco/endoplasmic reticulum Ca^2+^-ATPase; STIM1, Stromal interaction molecule 1; SK, intermediate-conductance calcium-activated potassium channels; TxA_2_, Thromboxane A_2_; TP, Thromboxane receptor. ↔ represents “no effect”, ↑ represents an “increase”. Created with BioRender.com.

## Author contributions

OO, SG conceived and designed the experiments; OO, SC conducted vaginal cytology examinations and animal mating protocols; OO conducted vascular reactivity, Western blot analyses, vascular bioassays and ELISA experiments; OO, SG drafted the manuscript; all authors edited and approved final version of the manuscript.

