## Supplemental Figures for "Cyclooxygenase-dependent mechanisms mediate in part the anti-dilatory effects of perivascular adipose tissue in uterine arteries from pregnant rats"

**Supplemental Figure 1:** Uncropped Western blot images of cyclooxygenase-1 (COX-1) and cyclooxygenase-2 (COX-2) in L-NAME pretreated uterine arteries incubated with or without PVATmedia. Representative images and statistical analysis are presented in Figure 7A-B.

**Supplementary Figure 2:** Relative protein expression of cyclooxygenase-1 (COX-1) and cyclooxygenase-2 (COX-2) in uterine arteries incubated with or without perivascular adipose tissue (PVAT)-conditioned media. Values are mean ± SEM. Paired t-test.

**Supplementary Figure 2:** Uncropped Western blot images of cyclooxygenase-1 (COX-1) and cyclooxygenase-2 (COX-2) in uterine arteries incubated with or without PVATmedia.

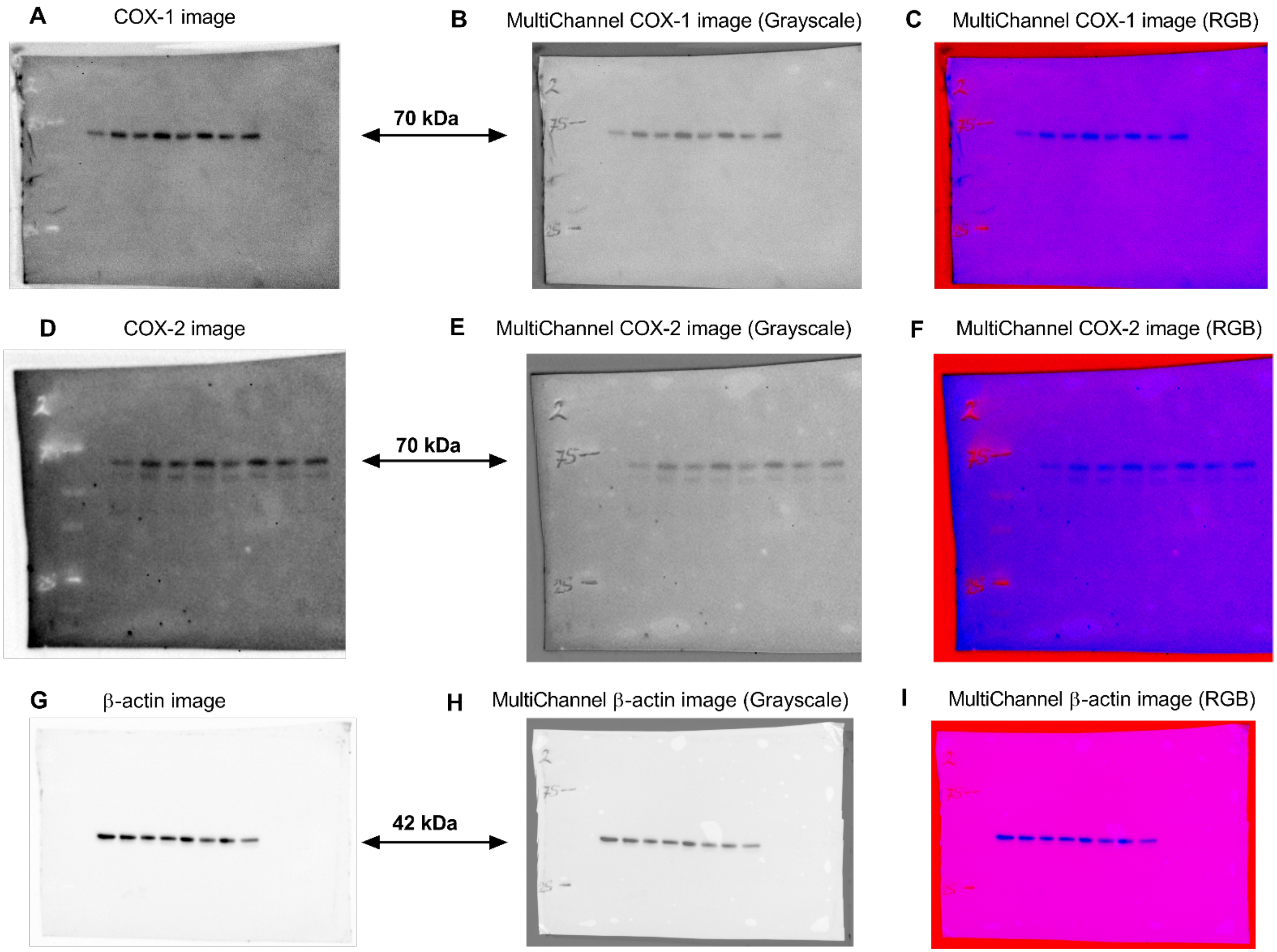
A
COX-1 image
B
MultiChannel COX-1 image (Grayscale)
C
MultiChannel COX-1 image (RGB)
70 kDa
D
COX-2 image
E
MultiChannel COX-2 image (Grayscale)
F
MultiChannel COX-2 image (RGB)
70 kDa
G
β-actin image
H
MultiChannel β-actin image (Grayscale)
I
MultiChannel β-actin image (RGB)
42 kDa

**Figure S1**

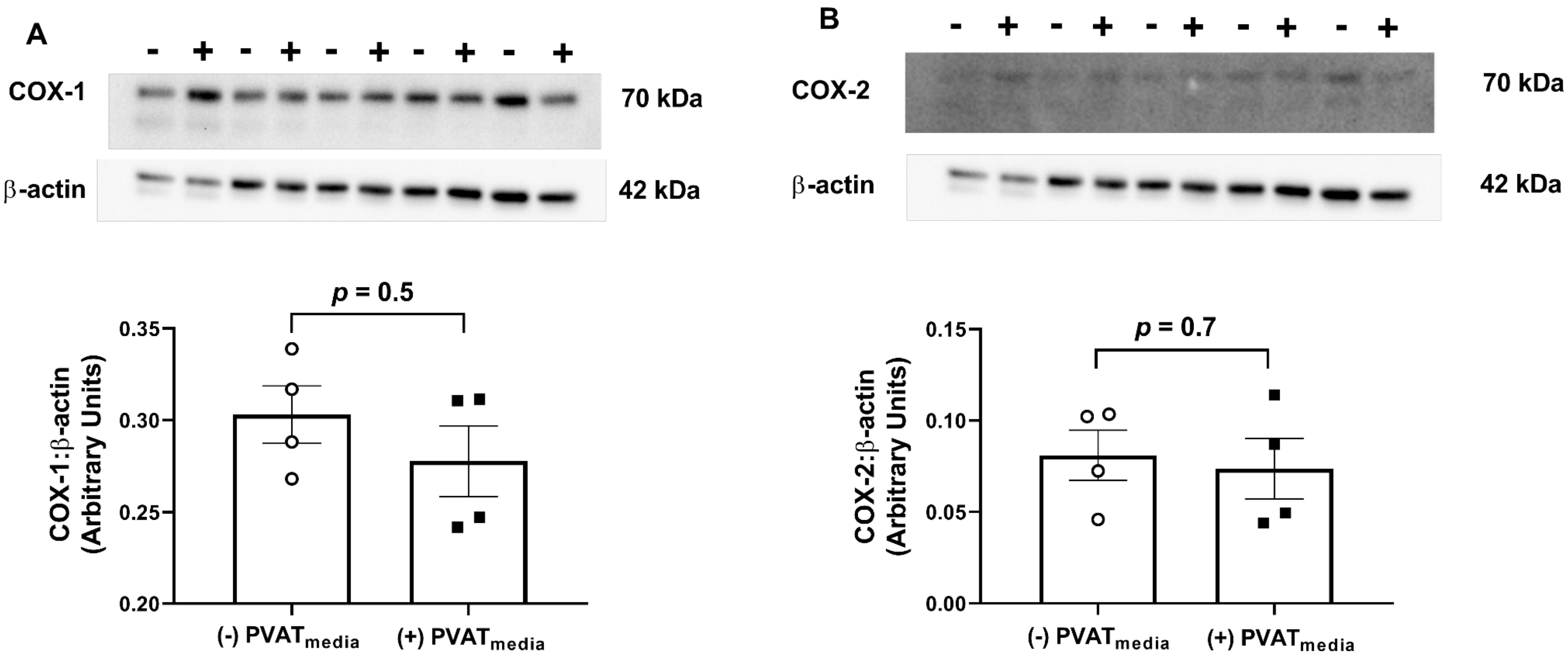
A
- + - + - + - + - +
COX-1
70 kDa
β-actin
42 kDa
p = 0.5
0.35
0.30
0.25
0.20
COX-1:β-actin (Arbitrary Units)
(-) PVATmedia
(+) PVATmedia
B
- + - + - + - + - +
COX-2
70 kDa
β-actin
42 kDa
p = 0.7
0.15
0.10
0.05
0.00
COX-2:β-actin (Arbitrary Units)
(-) PVATmedia
(+) PVATmedia

Figure S2

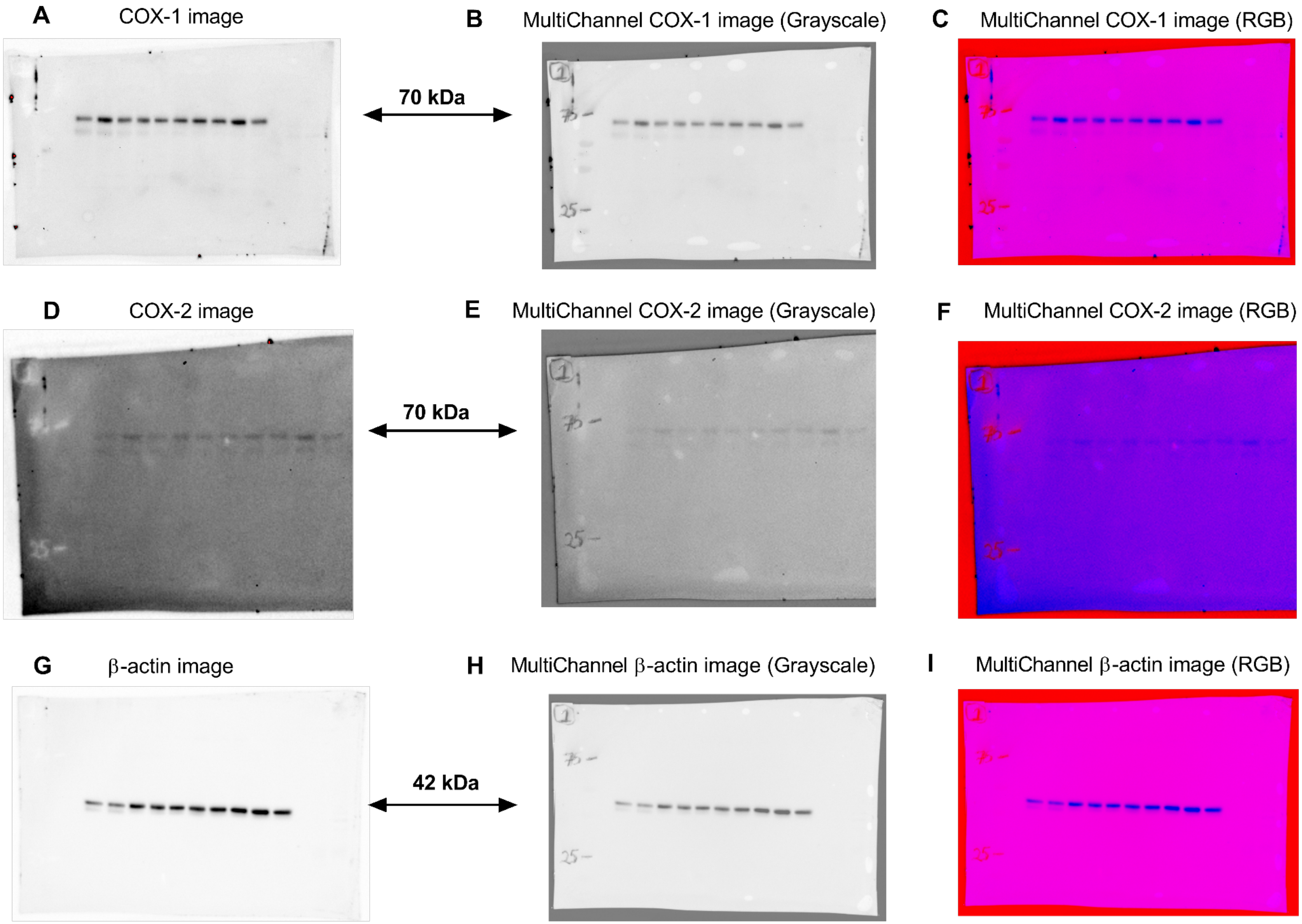
A
COX-1 image
B
MultiChannel COX-1 image (Grayscale)
C
MultiChannel COX-1 image (RGB)
70 kDa
D
COX-2 image
E
MultiChannel COX-2 image (Grayscale)
F
MultiChannel COX-2 image (RGB)
70 kDa
G
β-actin image
H
MultiChannel β-actin image (Grayscale)
I
MultiChannel β-actin image (RGB)
42 kDa

**Figure S3**
